# Reverse Transcription Loop-Mediated Isothermal Amplification Combined with Lateral Flow Assay (RT-LAMP-LFA) for Rapid Detection of the 2026 Bundibugyo Virus (BDBV) Outbreak Strain

**DOI:** 10.64898/2026.11.09.765523

**Authors:** Léo Shigueki Sato, Motoki Ihara, Arata Hidano, Yuki Takamatsu

## Abstract

Making accurate diagnosis is a critical first step in patient management and control for emerging and re-emerging disease outbreaks. Bundibugyo virus (BDBV) is a filovirus that causes Ebola disease, and the 2026 outbreak highlights the need for rapid diagnostic approaches that can be deployed in settings with limited laboratory infrastructure. Here, we developed a reverse transcription loop-mediated isothermal amplification–lateral flow assay (RT-LAMP-LFA) for rapid detection of the 2026 BDBV outbreak strain. LAMP primers were designed based on the genome sequence of the 2026 outbreak strain, and the conservation of the primer binding sites among all available 2026 outbreak genomes was confirmed in silico. LAMP primers were evaluated for analytical specificity and sensitivity using in vitro-transcribed RNA (ivtRNA) from BDBV and other filoviruses, as well as RNA derived from arboviruses and mammalian cells. Amplification products were detected using a Chromatography-Printed Array Strip (C-PAS) lateral flow platform. The newly designed primers specifically detected the presence of BDBV RNA without nonspecific amplification of the other viral or cellular RNA samples tested. Consistent amplification was observed down to 10^3^ copies per reaction. In addition, BDBV viral RNA was successfully amplified, and the amplification products were detected using C-PAS, with the total workflow being completed within 25 min. These findings demonstrate the analytical specificity and sensitivity of the developed RT-LAMP-LFA assay and its compatibility with rapid visual lateral flow detection, supporting its potential application as a point-of-care approach for BDBV detection.

**Importance:** Bundibugyo virus causes a form of Ebola disease for which rapid and reliable diagnostic testing is essential to support outbreak response. However, molecular tests may require specialized equipment and laboratory infrastructure, limiting access to testing in remote or resource-limited settings. In this study, we developed a rapid molecular assay targeting the 2026 outbreak strain of Bundibugyo virus. The assay detected viral RNA at concentrations as low as 10^3^ copies per reaction, showed no nonspecific amplification with the other viral and cellular RNA samples tested, and produced results using a visual detection platform within approximately 25 minutes, excluding sample preparation. These findings demonstrate the feasibility of a rapid and simplified approach for Bundibugyo virus detection. Further evaluation using clinical specimens and under field conditions is needed to establish its diagnostic performance and practical applicability.

## Introduction

On 17 May 2026, the World Health Organization (WHO) determined that the outbreak of Ebola disease caused by Bundibugyo virus (BDBV) in the Democratic Republic of the Congo (DRC) and Uganda constituted a Public Health Emergency of International Concern (PHEIC) [1]. BDBV is a member of the species *Orthoebolavirus bundibugyoense* within the genus *Orthoebolavirus* and family *Filoviridae* and is one of the causative agents of human Ebola disease [2,3]. Since its first identification during an outbreak in Uganda in 2007, BDBV has caused sporadic outbreaks, including one in the DRC in 2012 [4]. Effective control of Ebola disease requires rapid and accurate diagnosis, yet, unlike *Zaire ebolavirus* (EBOV), BDBV has no approved specific vaccines or therapeutics [5].

The early course of the 2026 BDBV outbreak illustrates the consequences of delayed diagnosis. The GeneXpert Ebola assay, widely used in the region for EBOV detection, failed to detect BDBV in initial samples [2,6]. Nearly three weeks elapsed between symptom onset in the presumed index case on 25 April and laboratory confirmation on 14 May. During this period, four healthcare workers died within four days. Diagnostic coverage has also remained limited: only 72% of validated alerts were tested, and only 15∼20% of new cases had a known infection source [7]. Limited understanding of transmission pathways, together with gaps in diagnostic capacity, may hinder timely case detection and containment, potentially facilitating further transmission. By 20 August, 158 healthcare workers had been infected and 45 had died, demonstrating intense transmission within healthcare settings [8]. In the absence of licensed BDBV-specific vaccines or therapeutics, early case identification and isolation remain the main control measures [6].

The WHO target product profile (TPP) for Ebola diagnostics, originally developed for EBOV [9,10], specifies desirable clinical sensitivity above 98% or acceptable sensitivity above 95%, together with analytical specificity above 99% [10,11]. These performance requirements reflect the serious consequences of both false-negative and false-positive results for individual patient management and public health responses. Although RT-qPCR remains the reference standard, five molecular tests currently hold WHO Emergency Use Listing for BDBV [12]. Nevertheless, these molecular diagnostic methods require specialized instrumentation, laboratory infrastructure, and trained personnel, hindering its use in resource-constrained and field settings [13]. These limitations are particularly relevant during viral hemorrhagic fever outbreaks, when diagnostic testing may be needed in decentralized healthcare facilities or areas with limited laboratory capacity [5]. Accordingly, rapid nucleic acid amplification tests that combine high diagnostic performance with simplified operation, such as loop-mediated isothermal amplification (LAMP) coupled with lateral flow detection (LAMP-LFA), are urgently required [14–19].

LAMP is a nucleic acid amplification method that operates under isothermal conditions and can be performed using relatively simple equipment. LAMP-based assays have been developed for filovirus detection, including EBOV and other known ebolavirus species [14–16]. However, previously reported BDBV assays were designed using sequence information available from earlier outbreaks [14–19]. Genetic differences between historical reference sequences and currently circulating strains may affect primer binding and amplification efficiency [17]. Furthermore, some existing assays require relatively high reaction temperatures, prolonged incubation, or additional equipment, which limit their use in resource-limited settings.

In the DRC, terrain, insecurity, and infrastructure constraints make even district-level healthcare facilities inaccessible to affected communities. In this context, this study aimed to develop an RT-LAMP-LFA assay for the rapid detection of the 2026 BDBV outbreak strain. The assay was evaluated for in silico conservation of the primer binding sites among currently circulating 2026 outbreak genomes, analytical specificity and sensitivity, detection in the presence of human plasma, and compatibility with viral RNA and the C-PAS lateral flow platform, with the aim of establishing a simple and rapid molecular detection approach suitable for field applications.

## Materials and methods

### In silico analysis of primer binding sites

To evaluate the conservation of the newly designed primer binding sites among currently circulating BDBV strains, the NP gene sequences of all complete BDBV genomes from the 2026 outbreak that were available at the time of analysis were aligned with representative NP gene sequences from the 2007 and 2012 outbreaks. Alignment was performed using MAFFT v7.5, and nucleotide mismatches within the binding regions of the six LAMP primers (eight regions: F3, F2, F1c, LF, B1c, LB, B2, and B3) were inspected. The number and position of mismatches, particularly those located at the 3′ ends of the primers, were evaluated. All sequences available at the time of analysis and included in the analysis are listed in Supplementary Table 1.

### Primer design

Primers targeting the NP gene were designed using the NEB^®^ LAMP Primer Design Tool (v1.5.1) [20] based on the genome sequence of the 2026 BDBV outbreak strain (PP_006XHKB) [21]. The designed primers were subsequently evaluated by in silico PCR using AmplifX2 (v2.1.1) [22]. The sequences of the primers designed in this study are shown in Table 1.

**Table 1.**
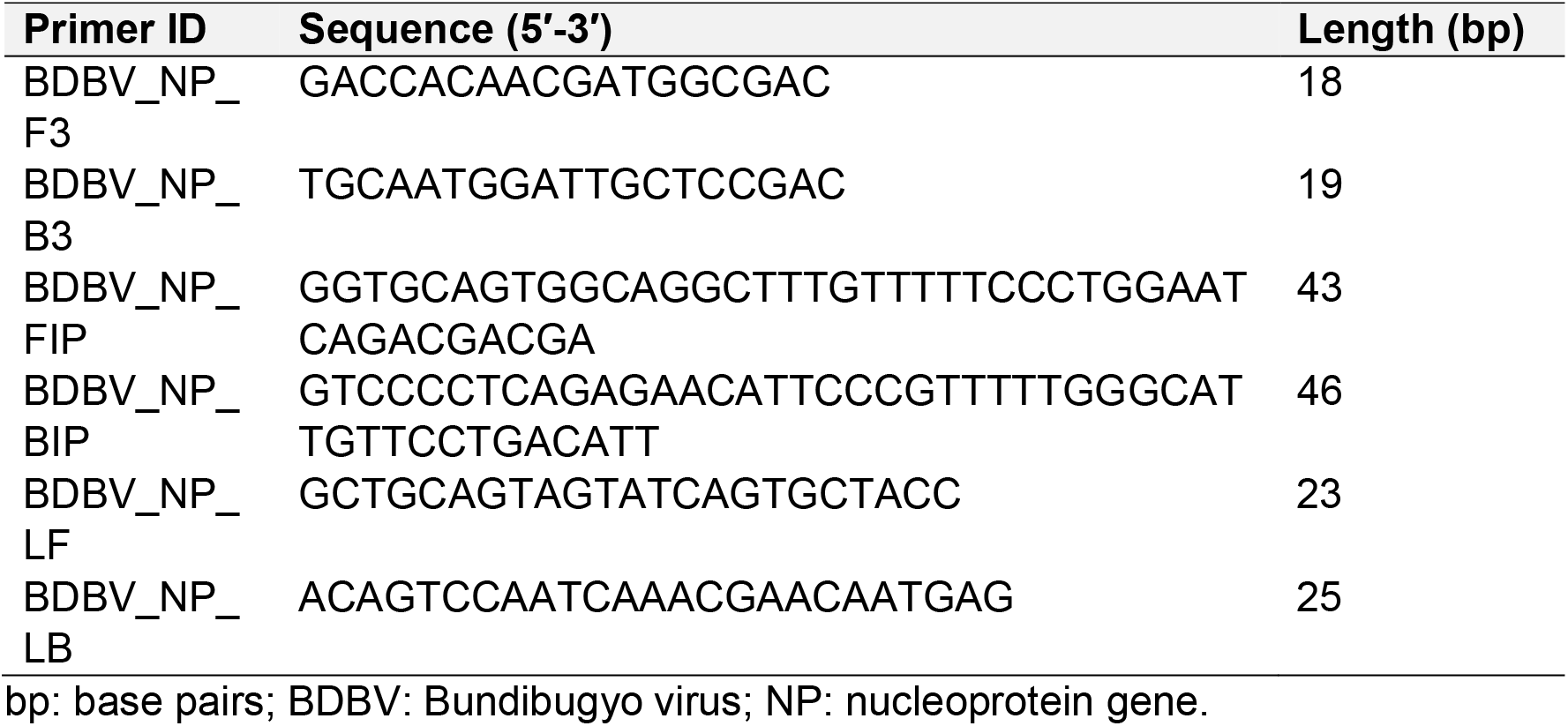
Sequences of the newly designed primers used in this study.

### Cells and viruses

The virus strains used in this study are listed in Table 2. Full-length NP gene DNA fragments of BDBV (2,959 bp), Bombali virus (BOMV; 2,964 bp), Sudan virus (SUDV; 2,952 bp), Reston virus (RESTV; 2,958 bp), and Taï Forest virus (TAFV; 2,965 bp) were purchased from GenScript Japan. DNA fragments of EBOV and Marburg virus (MARV) were obtained as previously described [23]. Dengue virus (DENV) and Yellow Fever virus (YFV) were provided by Nagasaki University through the National BioResource Project (NBRP; Human Pathogenic Viruses) of the Ministry of Education, Culture, Sports, Science and Technology (MEXT), Japan [24]. Zika virus (ZIKV) and Chikungunya virus (CHIKV) were kindly provided by Dr. Mya Myat Ngwe Tun (Institute of Tropical Medicine, Nagasaki University, Japan). HEK-293T cells were purchased from the American Type Culture Collection (ATCC; CRL-3216), and Vero cells from KAC Co., Ltd. (IU-002).

**Table 2.**
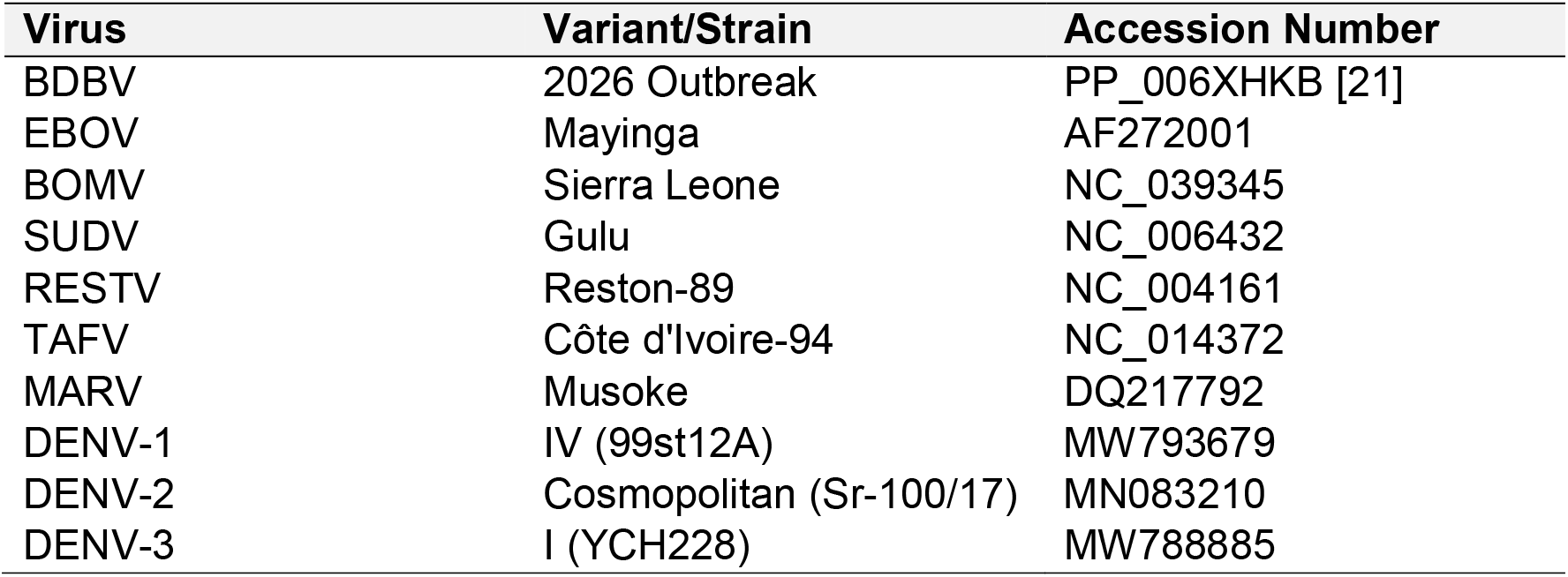
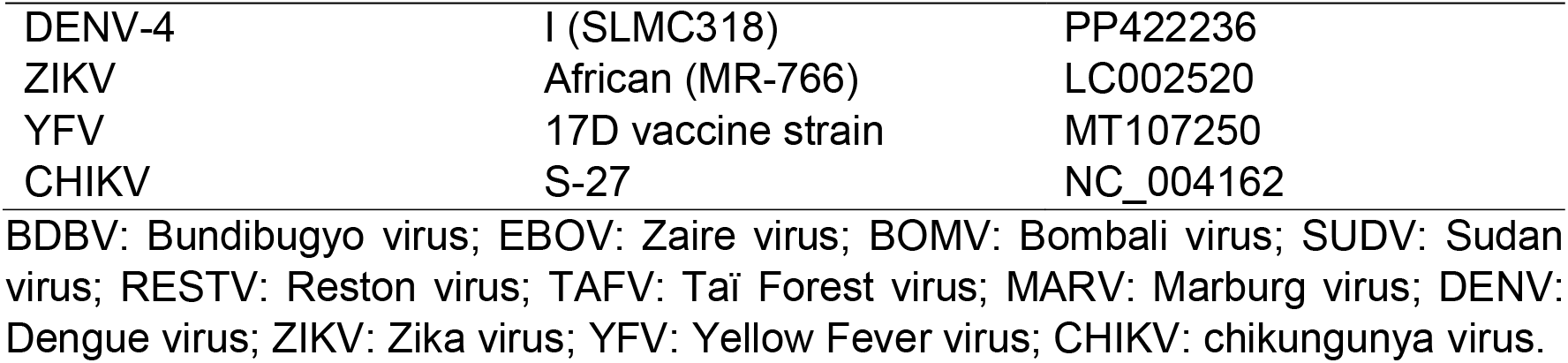
Viruses used to determine the specificity and sensitivity of RT-LAMP.

### Viral and cellular RNA extraction

Viral RNA extraction was performed using the QIAamp^®^ Viral RNA Kit (QIAGEN, Hilden, Germany) according to the manufacturer’s instructions. Total RNA from mammalian cells was extracted using ISOGEN II (Nippon Gene, Toyama, Japan) following the manufacturer’s instructions. All the extracted RNA was stored at −80 °C until further use.

### Plasmid DNA production and artificial RNA synthesis

Full-length NP gene DNA fragments of BDBV, EBOV, BOMV, SUDV, RESTV, TAFV, and MARV were cloned using the Zero Blunt^®^ TOPO^®^ PCR Cloning Kit (Thermo Fisher Scientific, MA, USA) according to the manufacturer’s instructions, generating plasmids containing the T7 promoter. Plasmid DNA was subsequently purified using the QIAprep^®^ Spin Miniprep Kit (QIAGEN) according to the manufacturer’s instructions.

Using the purified plasmid DNA as a template, RNA transcripts were synthesized by in vitro transcription followed by DNase treatment using an In-vitro Transcription T7 Kit (TaKaRa Bio, Shiga, Japan), according to the manufacturer’s instructions. The transcripts were purified using the QIAamp^®^ Viral RNA Kit (QIAGEN) according to the manufacturer’s instructions. In vitro transcribed RNA (ivtRNA) concentration and purity were determined using a NanoDrop™ ND-1000 spectrophotometer (Thermo Fisher Scientific).

### One-step RT-LAMP

One-step RT-LAMP reaction was performed using the Isothermal Amplification Buffer II Pack (New England Biolabs, MA, USA) in a total volume of 25 µL containing 2.5 µL of 10x Isothermal Amplification Buffer II, 0.5 µL of MgSO4 (100 mM), 3.5 µL of dNTP mix (10 mM), 1.6 µM each of the FIP and BIP primers, 0.8 µM each of the LF and LB primers, 0.2 µM each of the F3 and B3 primers, 1 µL of Bst 3.0^®^ DNA Polymerase (8,000 U/mL), and 5 µL of RNA template (approximately 50 ng). Reverse transcription of the RNA template and subsequent isothermal amplification were performed relying on the reverse transcription activity of Bst 3.0 DNA polymerase, without the addition of a separate reverse transcriptase. The reaction was incubated at 60 °C for 20 min using an Applied Biosystems™ SimpliAmp™ Thermal Cycler (Thermo Fisher Scientific). The reaction temperature was optimized by evaluating

LAMP amplification at different incubation temperatures. The incubation time was subsequently optimized under the selected temperature condition by assessing amplification at different reaction times. Amplification was assessed by adding 2 µL of 1:1000 diluted SYBR^®^ Green I Nucleic Acid Stain (Lonza, Basel, Switzerland) and visualizing under blue LED illumination. RT-LAMP products were further analyzed by electrophoresis on a 2% agarose gel at 100 V for 30 min and visualized under blue LED illumination.

### Specificity and sensitivity analysis

The analytical specificity of the newly designed LAMP primer sets was evaluated using ivtRNA from BDBV, EBOV, BOMV, SUDV, RESTV, TAFV, and MARV. In addition to ivtRNA from filoviruses, viral RNA extracted from DENV-1–4, ZIKV, YFV, and CHIKV, as well as cellular RNA extracted from HEK-293T and Vero cells, was included in the specificity analysis. Each RNA sample was tested at 50 ng per reaction.

The analytical sensitivity of the newly designed LAMP primer sets was evaluated using ten-fold serial dilutions of BDBV ivtRNA, ranging from 10^7^ to 10^0^ copies per reaction. The RNA standard was generated by in vitro transcription of the BDBV NP DNA fragment. RNA concentration was determined using a Qubit^®^ 2.0 Fluorometer and Qubit™ RNA HS (High Sensitivity) Assay Kit (Thermo Fisher Scientific) according to the manufacturer’s instructions, and RNA copy number was calculated based on the molecular weight and length of the transcribed RNA fragment. The RNA standard was subsequently quantified by RT-qPCR using the primers and probe listed in Table 3, with each standard concentration analyzed in duplicate.

**Table 3.**
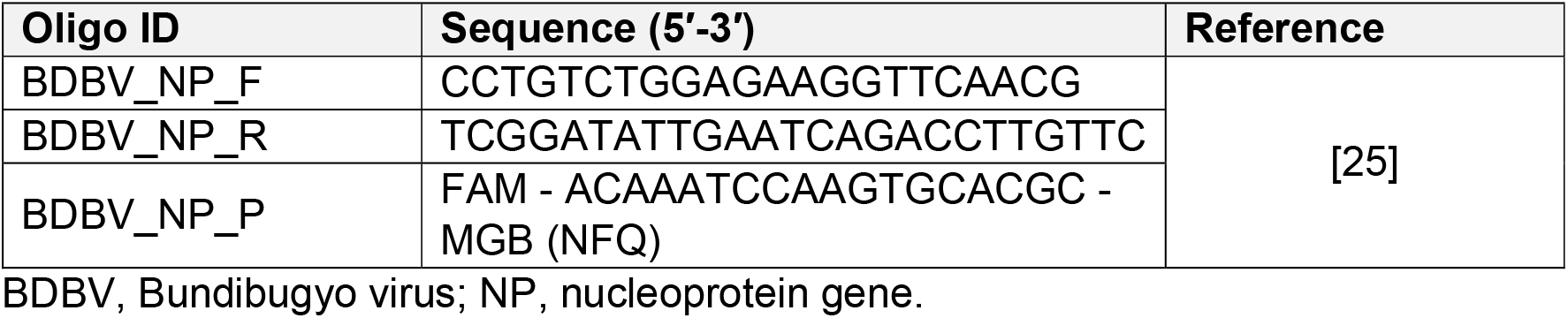
Primers and probe used for BDBV RT-qPCR.

The BDBV RT-qPCR assay was performed using the Applied Biosystems™ TaqMan™ Fast Virus 1-Step Master Mix (Thermo Fisher Scientific) in a final reaction volume of 25 µL containing 6.25 µL of 4x TaqMan™ Fast Virus 1-Step Master Mix, 0.3 µM of forward primer, 0.9 µM of reverse primer, 0.2 µM of probe, and 2 µL of template RNA. This protocol was adapted from the European Union Reference Laboratory (EURL) [25], with modification in template RNA volume. Amplification and fluorescence detection were performed using an Applied Biosystems™ QuantStudio™ 5 Real-Time PCR System (Thermo Fisher Scientific), and the data were analyzed using QuantStudio™ Design & Analysis Software v2.8.0 (Thermo Fisher Scientific).

### Detection in in vitro-spiked human samples

The RT-LAMP assay was also evaluated using human plasma spiked with BDBV ivtRNA to assess whether plasma components interfered with the amplification reaction. Plasma collected from a healthy individual was spiked directly with BDBV ivtRNA at 10^4^ copies per reaction, and the spiked plasma was added to the reaction mix directly. The same plasma sample without added RNA was used as the negative control (non-spiked plasma).

### BDBV viral RNA detection and lateral flow assay (LFA)

The newly designed LAMP primers were tested using BDBV viral RNA, which was kindly provided by Dr. Kim Heimsch and Prof. Christian Drosten from Charité – Universitätsmedizin Berlin and the Emergency Supply Coordination Team of WHO. The BDBV viral RNA was detected using a combined LAMP-lateral flow assay (RT-LAMP-LFA) with Chromatography-Printed Array Strip (C-PAS) (Tohoku Bio-Array [TBA] Co., Ltd., Miyagi, Japan) following a 5 min incubation. For the RT-LAMP-LFA assay, the LF primer was 5′-biotin-labelled, whereas the LB primer was modified with a proprietary tag sequence provided by the manufacturer (TBA Co., Ltd.).

## Results

### In silico conservation of the primer binding sites

The newly designed LAMP primer set targets the NP gene of BDBV and comprises six primers, including the outer primers F3 and B3, the inner primers FIP (F2/F1c) and BIP (B1c/B2), and the loop primers LF and LB (Figure 1A). We assessed the conservation of the primer-binding sites by comparing the NP gene sequence of the 2026 BDBV outbreak strain (PP_006XHKB) used for primer design with representative BDBV genomes from the 2007, 2012, and 2026 outbreaks. Compared with the BDBV reference sequence (NC_014373), one nucleotide mismatch was identified in the F3 region and one in the B2 region of BIP. In addition, a single nucleotide mismatch in the B1c region of BIP was consistently observed among the 2012 outbreak strains included in the analysis (Figure 1B). The geographic and temporal distribution of variation at the B1c-binding site showed that this mismatch was present in the analyzed 2012 outbreak sequences, whereas the site was conserved among the complete BDBV genomes from the 2026 outbreak available at the time of analysis (Figure 1C). All six primer-binding regions were fully conserved among these 2026 outbreak genomes, including the strain used for primer design. These results indicate that the designed LAMP primers target highly conserved regions of the NP gene among the BDBV sequences analyzed from the 2026 outbreak.

**Figure 1.**
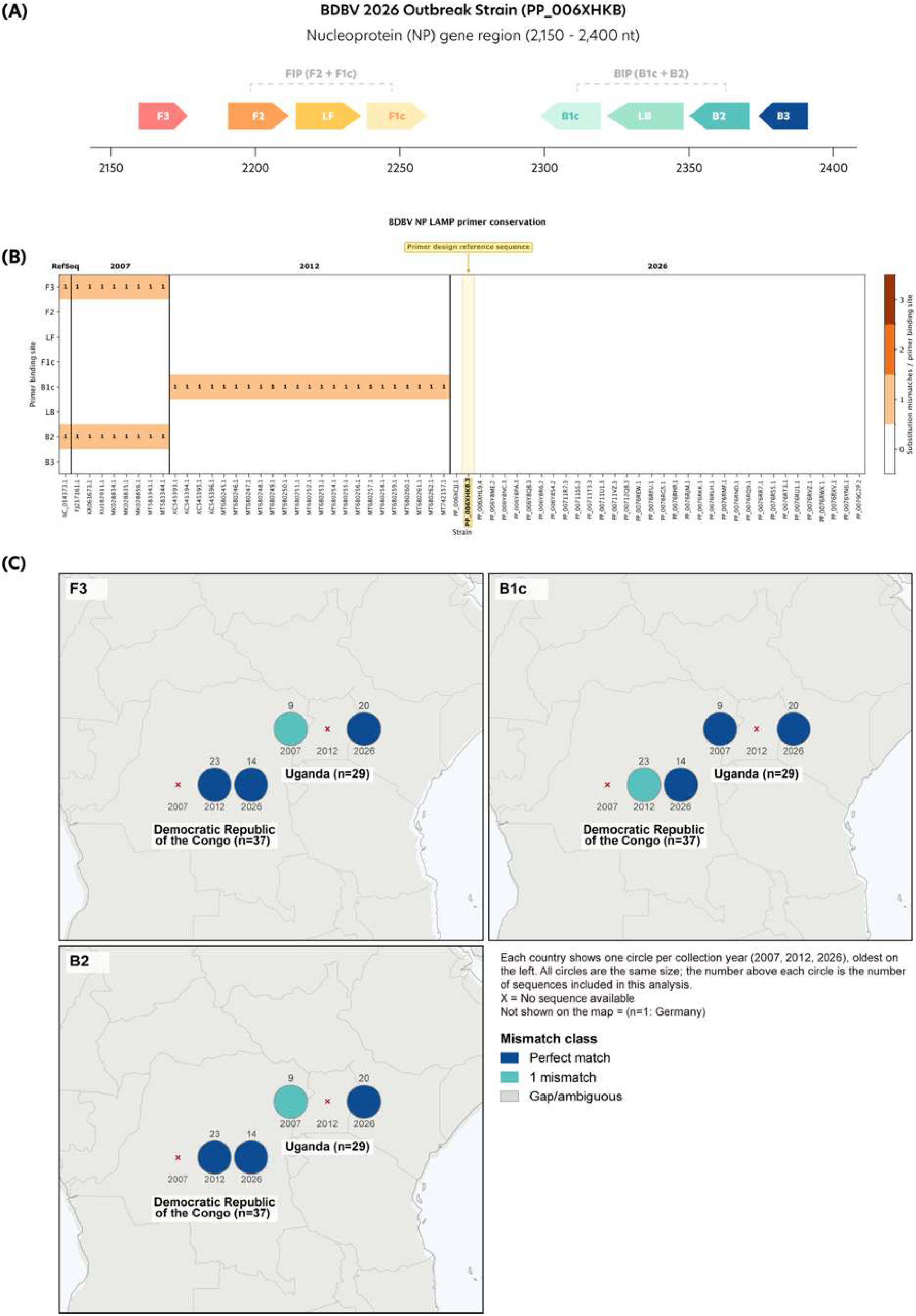
BDBV NP gene target regions and conservation of the LAMP primer-binding sites. (A) Schematic representation of the BDBV NP gene region targeted by the LAMP primers. FIP comprises the F2 and F1c regions, and BIP comprises the B1c and B2 regions. (B) Sequence conservation of the primer-binding sites among BDBV strains from the 2007, 2012, and 2026 outbreaks. Numbers indicate the number of nucleotide mismatches between the primer-binding sites and the corresponding viral sequences. The 2026 outbreak sequence used as the primer-design reference is indicated by the yellow marker. (C) Geographic and temporal distribution of sequence variation at BDBV RT-LAMP primer-binding sites. The maps show the distribution of sequences from the DRC and Uganda across the 2007, 2012, and 2026 outbreaks, with sequence variation at the binding sites of the F3, B1c, and B2 primer-binding sites. X = No sequence available. RefSeq, reference sequence; FIP, forward inner primer; BIP, backward inner primer; LF, loop-forward primer; LB, loop-back primer.

### Specificity analysis

IvtRNA of BDBV, EBOV, BOMV, SUDV, RESTV, TAFV, and MARV, as well as viral RNA extracted from DENV-1–4, ZIKV, YFV, and CHIKV and cellular RNA extracted from HEK-293T and Vero cells, was included in the analytical specificity assessment of the newly designed LAMP primer sets. The primer sets specifically detected the in vitro transcribed BDBV RNA, with no nonspecific amplification observed for any of the other viral or cellular RNA samples tested. SYBR^®^ Green detection under blue LED illumination showed strong fluorescence exclusively in the reaction containing BDBV RNA (Figure 2A). Consistently, agarose gel electrophoresis revealed the characteristic LAMP amplification smear only for the BDBV RNA reaction, further supporting the specificity of the newly designed primer sets (Figure 2B).

**Figure 2.**
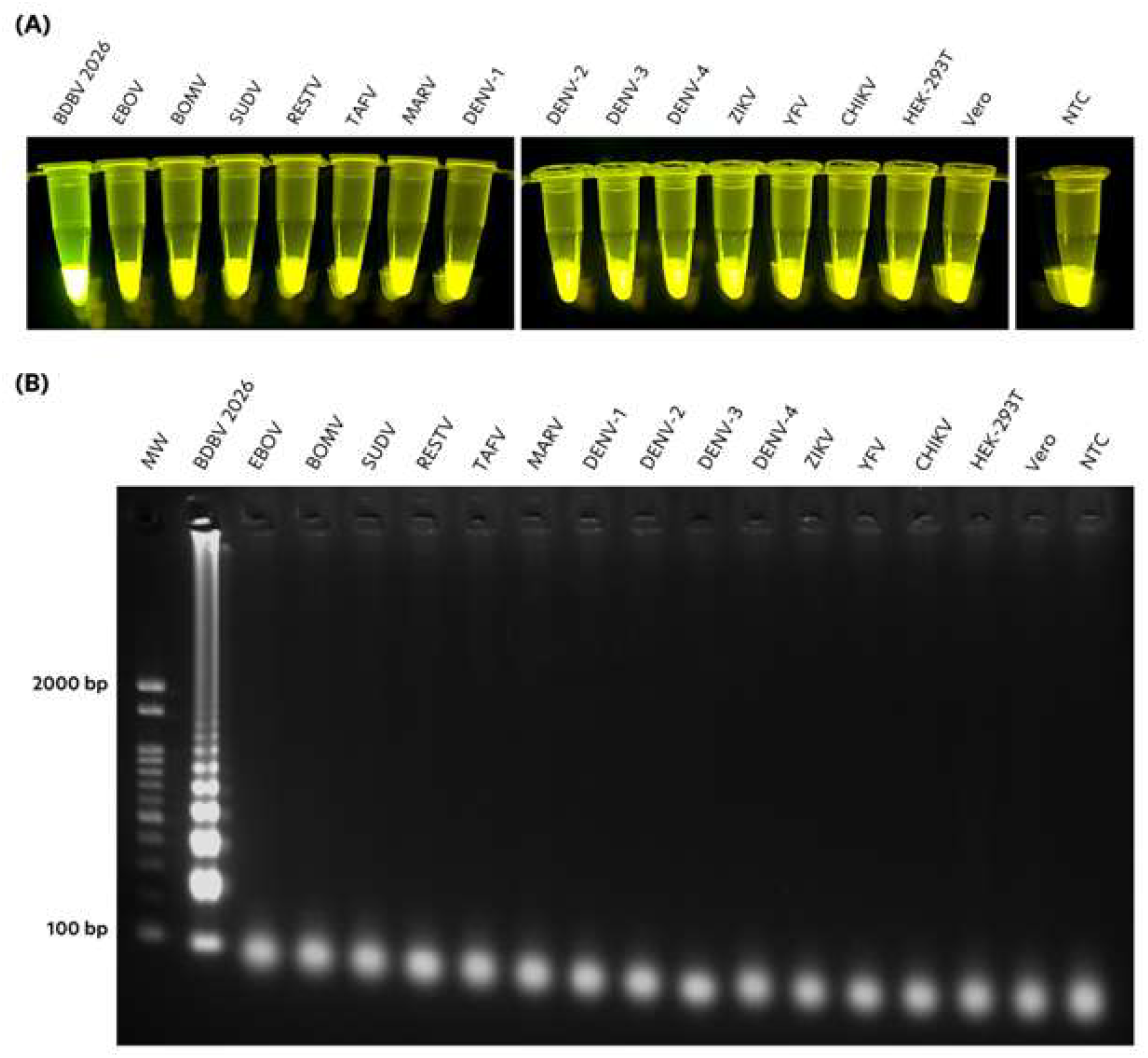
Analytical specificity of the newly designed LAMP primer sets for BDBV detection. In addition to ivtRNA from the 2026 BDBV outbreak strain, ivtRNA from other filoviruses (EBOV, BOMV, SUDV, RESTV, TAFV, and MARV), arboviruses (DENV-1–4, ZIKV, YFV, and CHIKV), and mammalian cell lines (HEK-293T and Vero) was included to evaluate primer specificity. The primer sets specifically detected BDBV ivtRNA, with no nonspecific amplification observed for any of the other viral or cellular RNA samples tested. (A) SYBR^®^ Green fluorescence visualized under blue LED illumination. (B) Electrophoretic analysis on a 2% agarose gel. MW, molecular weight; kb, kilobases; BDBV, Bundibugyo virus; EBOV, Ebola virus; BOMV, Bombali virus; SUDV, Sudan virus; RESTV, Reston virus; TAFV, Taï Forest virus; DENV, dengue virus; ZIKV, Zika virus; YFV, yellow fever virus; CHIKV, chikungunya virus; HEK-293T, human embryonic kidney 293T cells; Vero, African green monkey kidney cells; NTC, non-template control; bp, base pairs.

### Sensitivity analysis

The analytical sensitivity of the newly designed primer set was evaluated using ten-fold serial dilutions of BDBV ivtRNA, from 10^7^ to 10^0^ copies per reaction. The primers detected the RNA at low concentrations, with consistent amplification down to 10^3^ copies per reaction (Figure 3). All the experiments were carried out in duplicate.

**Figure 3.**
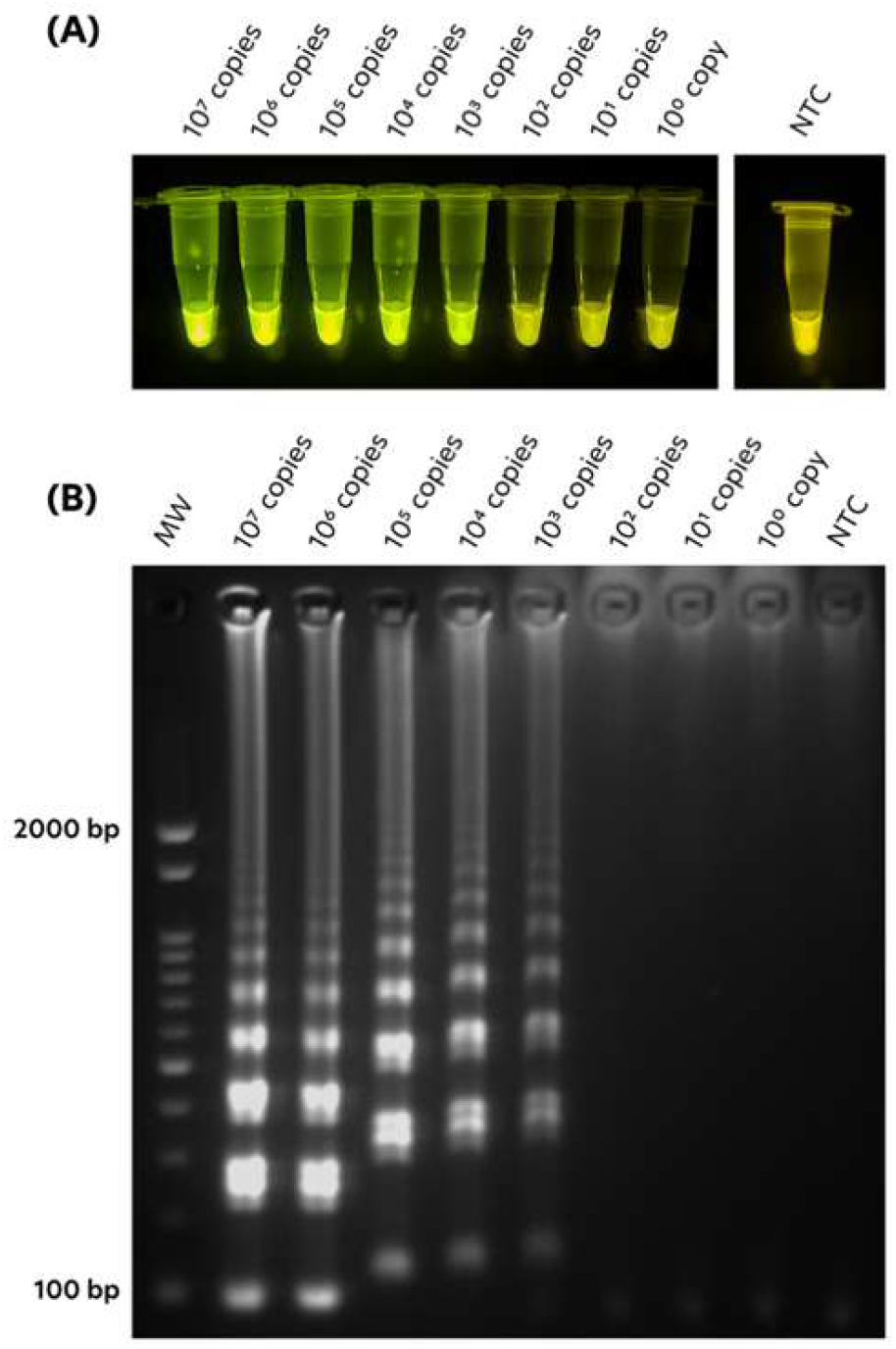
Analytical sensitivity of the newly designed LAMP primer set. The sensitivity was evaluated using 10-fold serial dilutions, starting from an initial concentration of 10^7^ copies/reaction of BDBV RNA standard. The primers detected and amplified the BDBV NP gene at low concentrations, with consistent amplification down to 10^3^ copies per reaction. (A) SYBR^®^ Green fluorescence visualized under blue LED illumination. (B) Electrophoretic analysis on a 2% agarose gel. MW, molecular weight; NTC, non-template control; bp, base pairs.

### Detection in in vitro-spiked human plasma

The newly designed LAMP primer sets were evaluated using human plasma samples spiked with BDBV ivtRNA to assess potential interference from plasma components (Figure 4). The primer sets detected the BDBV ivtRNA in the presence of human plasma, indicating that plasma components did not interfere with the LAMP reaction. These results support the potential applicability of the newly designed primer sets for BDBV detection in clinical samples.

**Figure 4.**
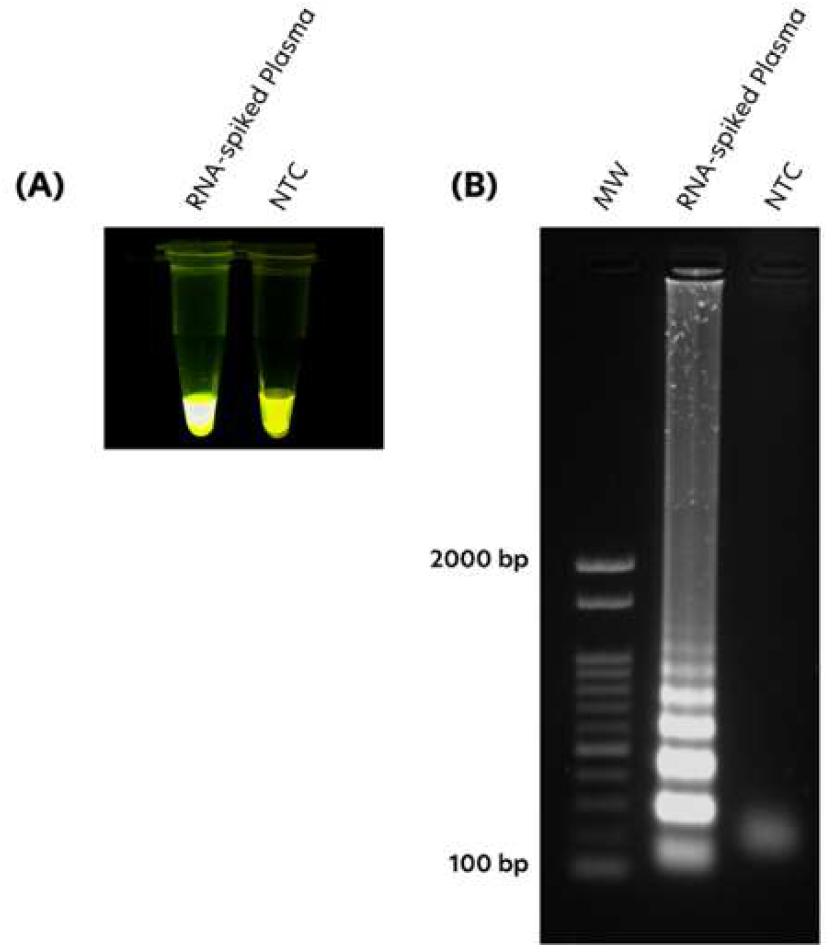
Evaluation of the BDBV RT-LAMP assay using ivtRNA-spiked human plasma. BDBV ivtRNA was spiked directly into plasma collected from a healthy individual to evaluate whether plasma components interfered with amplification. The same plasma sample without added RNA was used as the negative control (non-spiked plasma). RT-LAMP products were analyzed by (A) SYBR^®^ Green I visualization under blue LED illumination and (B) electrophoresis on a 2% agarose gel. Amplification was observed only in the RNA-spiked plasma sample, indicating that plasma components did not inhibit amplification or produce nonspecific amplification. MW, molecular weight; kb, kilobases; NTC, non-template control (non-spiked plasma).

### BDBV viral RNA detection and RT-LAMP-LFA reaction

BDBV viral RNA was also evaluated as a template for RT-LAMP amplification. The BDBV viral RNA was kindly provided by Dr. Kim Heimsch from Charité – Universitätsmedizin Berlin and the Emergency Supply Coordination Team of the WHO. Following RT-LAMP amplification, the reaction products were applied to the C-PAS and evaluated after the recommended incubation period. As shown in Figure 5, the BDBV-specific test line was clearly detected, indicating successful amplification of the target RNA. The flow control line was also observed during the assay and disappeared after complete migration of the reaction mixture through the strip, confirming appropriate sample flow and valid test performance.

**Figure 5.**
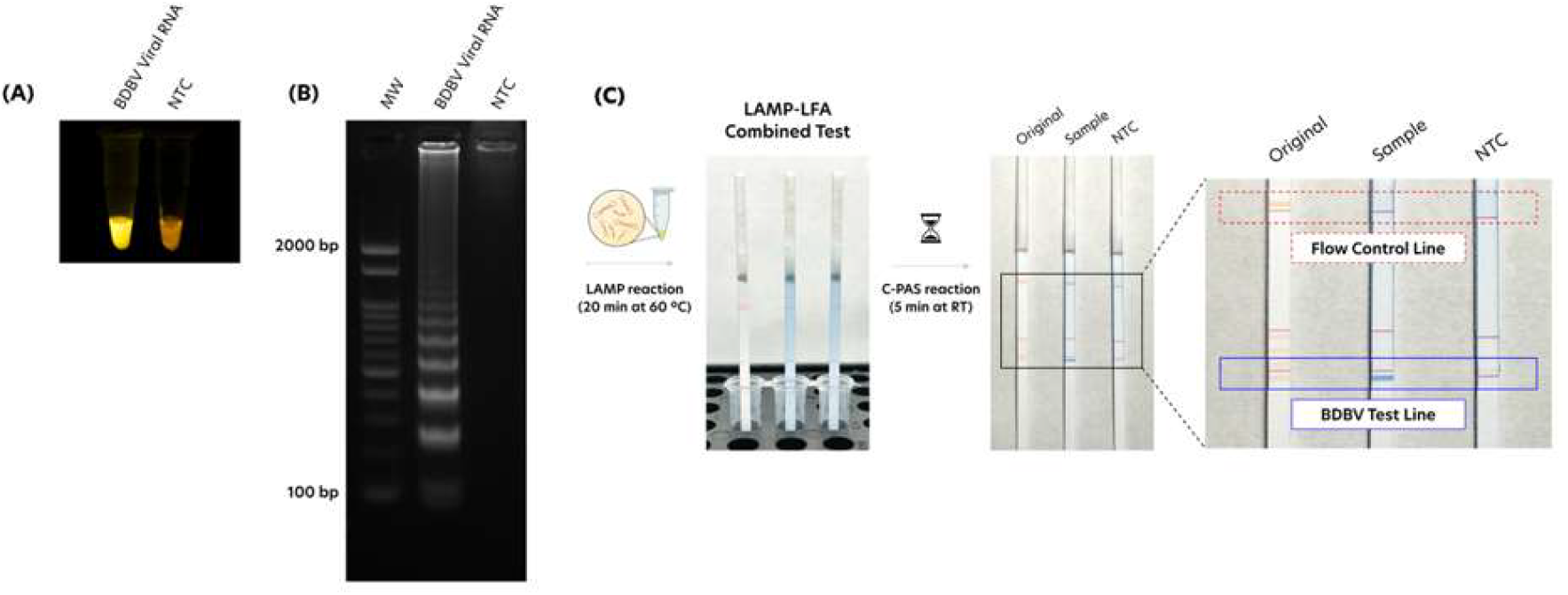
Evaluation of the BDBV RT-LAMP assay using BDBV viral RNA. RT-LAMP products were analyzed by (A) SYBR^®^ Green I visualization under blue LED illumination and (B) electrophoresis on a 2% agarose gel. (C) Workflow and representative results of lateral flow detection using the C-PAS device. Following RT-LAMP amplification, the C-PAS strip was applied to the reaction products, and the results were interpreted after the recommended incubation period. The enlarged image highlights the flow control line (red box), which appears during chromatography and disappears once the reaction mixture has completely migrated through the strip, confirming proper sample flow and valid test performance, and the BDBV-specific test line (blue box), indicating successful amplification of the target RNA (Original: non-reacted strip before application to the LAMP product; Sample: BDBV viral RNA; NTC: non-template control). BDBV, Bundibugyo virus; LAMP, loop-mediated isothermal amplification; LFA, lateral flow assay; NP, nucleoprotein gene; C-PAS, chromatography-printed array strip.

## Discussion

Using BDBV genome sequenced during the 2026 outbreak, we identified target regions that remain conserved across the geographic range of the current epidemic and developed an RT-LAMP-LFA assay detecting 10^3^ copies/reaction. Existing BDBV-detecting assays were designed against the 2007 and 2012 outbreak sequences, and their inclusivity against currently circulating genomes has not been reported. In this context, our assay resolves this gap by anchoring primer design to contemporaneous sequence data.

In the present study, all eight primer-binding regions were fully conserved among the complete 2026 outbreak BDBV genomes available at the time of analysis. A critical consideration in primer design was the genomic divergence of the 2026 outbreak strain from the BDBV prototype sequence (NC_014373). A previous study identified 219 nucleotide differences between the 2026 outbreak strain and the 2007– 2008 Uganda strain, as well as 216–227 nucleotide differences between the 2026 outbreak strain and the 2012 DRC strains, corresponding to approximately 1.2% sequence divergence across the genome [26]. This divergence is large enough to affect the performance of diagnostic assays designed using historical reference sequences. The conservation of all eight primer-binding regions supports the suitability of the selected target regions for the strains represented in this dataset. Nevertheless, continued assessment of newly available sequences will be important to monitor emerging variation in primer-binding regions and assess its potential impact on assay performance. The advantage of our platform is that we can tailor the assay to future ebolavirus strains once sufficient sequence information is accumulated.

The analytical specificity of the assay was supported by the absence of detectable nonspecific amplification with the other filoviruses, arboviruses, and mammalian cell-derived RNA included in the evaluation. This is relevant in the DRC, where filoviruses have previously caused outbreaks [27] and several of the arboviruses evaluated in this study are endemic [28]. As the consequence of false-positive diagnosis is devastating, further validation using clinical specimens from the DRC is required to quantify its diagnostic specificity.

The assay detected BDBV ivtRNA at concentrations as low as 10^3^ copies per reaction. In addition, the assay was able to detect and amplify BDBV ivtRNA spiked directly into human plasma at 10^4^ copies per reaction despite the presence of plasma components that may interfere with nucleic acid amplification, including myoglobin, EDTA, and immunoglobulins [29,30]. These findings are encouraging when considered alongside evidence of high viremia during acute filovirus infection: in a patient who recovered from acute BDBV infection, plasma viral RNA peaked at approximately 7 × 10^6^ copies/mL on day 5 of illness [31]. In patients with Ebola virus disease, viremia can reach approximately 10^9^ copies/mL, peaking around days 5–7 after symptom onset [32]. For context, 5 µL of plasma containing 10^7^ copies/mL is equal to approximately 5 × 10^4^ copies of viral RNA, well above the detection limit observed here, even if used directly as the reaction template.

Importantly, BDBV viral RNA was also successfully amplified and detected using the C-PAS lateral flow platform, demonstrating the feasibility of the complete RT-LAMP-LFA workflow. The protocol required 20 min for RT-LAMP at 60 °C, followed by 5 min for lateral flow detection at room temperature, yielding a nominal analytical turnaround time of approximately 25 min, excluding sample preparation. Rather than replacing real-time RT-PCR-based confirmation at reference laboratories, the assay could serve as a presumptive first-line test at peripheral facilities, enabling earlier isolation of suspected cases while confirmatory testing is arranged. The short reaction time and visual readout may be advantageous in outbreak settings where access to conventional molecular diagnostic platforms is limited. Furthermore, rapid on-site testing could reduce the need to transport specimens from collection sites to centralized diagnostic facilities, potentially shortening the time to obtain results. During the early phase of the outbreak, specimen collection, transportation, and diagnostic testing reportedly took approximately 3–4 days [33]. Nevertheless, this potential advantage remains to be confirmed through field evaluation, including assessment of sample preparation requirements and the assay’s performance under actual outbreak conditions.

Several limitations of this study should be acknowledged. First, analytical sensitivity was evaluated using synthetic RNA, with each dilution tested in duplicate; the lowest concentration at which amplification was consistently observed was 10^3^ copies per reaction. In addition, the LFA kit used here remains under development as part of an integrated LAMP–LFA format, and the number of validation runs performed to date is limited. Second, the plasma experiments were conducted using spiked samples rather than clinical specimens from BDBV-infected patients. Evaluation using well-characterized clinical specimens, including samples from patients infected with BDBV, will be essential to determine the assay’s diagnostic performance. Third, the open-tube handling required after LAMP amplification may increase the risk of carry-over contamination through the generation of aerosolized amplicons [34]. Developing a fully closed detection system will therefore be an important step toward reducing contamination risk and advancing the assay for point-of-care applications. To this end, an integrated LAMP–LFA kit using lyophilised reagents, which would require only a simple heat block for detection, is currently being developed in collaboration with an industrial partner. Finally, RNA extraction relied on commercially available kits requiring laboratory equipment, including a centrifuge, which may limit the assay’s applicability in resource-limited settings. Preliminary results indicate that the assay can partially amplify target RNA directly without prior RNA extraction; we are therefore evaluating a simplified sample-preparation workflow together with assay robustness under variable environmental conditions (e.g. fluctuations in temperature and humidity), and aim to further improve its analytical accuracy.

In conclusion, the RT-LAMP-LFA assay developed in this study provides a rapid approach for detecting the 2026 BDBV outbreak strain, combining analytical specificity against the tested panel, detection of BDBV RNA at low copy numbers, and visual readout using the C-PAS lateral flow platform. The observed analytical detection levels, considered alongside reported viral loads during acute BDBV infection, support further investigation of the assay’s potential for clinical diagnosis. Further validation using well-characterized patient specimens and development of a practical sample-preparation workflow will be essential to establish its diagnostic performance and advance its potential application in BDBV diagnosis, surveillance, and outbreak response.

## Authors’ contributions

Conceptualization: [YT]; Data curation: [LSS, MI, AH, YT]; Formal analysis: [LSS*, MI*]; Funding acquisition: [YT]; Investigation: [LSS*, MI*]; Methodology: [LSS*, MI*, YT]; Project administration: [LSS*, MI*]; Resources: [YT]; Software: [LSS, MI]; Supervision: [AH, YT]; Validation: [LSS*, MI*, YT]; Visualization: [LSS*, MI*]; Writing – original draft: [LSS, YT]; Writing – review & editing: [LSS, MI, AH, YT]. All authors read and approved the final manuscript.

*These authors contributed equally to this work.

## Acknowledgments

The authors are grateful to Prof. Christian Drosten and Dr. Kim Heimsch from Charité - Universitätsmedizin Berlin and the Emergency Supply Coordination Team of WHO for their kind gift of BDBV viral RNA. The authors also thank the teams at the Institut National de Recherche Biomédicale (INRB, Democratic Republic of the Congo) and the Central Public Health Laboratory (CPHL, Uganda), including Prof. Placide Mbala-Kingebeni and Dr. Isaac Ssewanyana, and their collaborators, for making the genome sequences of the 2026 BDBV outbreak strains available through Pathoplexus, and the virological.org community for rapid genomic data sharing. The authors also wish to thank all members of the Department of Virology, Institute of Tropical Medicine (NEKKEN), Nagasaki University. L.S.S. acknowledges the Japan International Cooperation Agency (JICA) for the scholarship support provided through the “Scholarship for Nikkei Communities in Latin America and the Caribbean: Program for Developing Leaders in Nikkei Communities.”

## Data availability statement

The newly designed LAMP primer sequences are provided in Table 1. The 2026 BDBV outbreak genome sequence used for primer design is publicly available from Pathoplexus (https://pathoplexus.org/seq/PP_006XHKB). All sequences available at the time of analysis and included in the analysis are listed in Supplementary Table 1. All raw gel and lateral flow images generated in this study are available from the corresponding author upon reasonable request.

## Declaration of generative AI use

Generative AI tools were used for language editing and preparation of this manuscript. The authors reviewed and edited all content generated with the assistance of AI tools and take full responsibility for the accuracy and integrity of the published work.

